# Expansion Microscopy provides new insights into the cytoskeleton of malaria parasites including the conservation of a conoid

**DOI:** 10.1101/2020.07.08.192328

**Authors:** Eloïse Bertiaux, Aurélia C Balestra, Lorène Bournonville, Mathieu Brochet, Paul Guichard, Virginie Hamel

## Abstract

Malaria is caused by unicellular *Plasmodium* parasites. *Plasmodium* relies on diverse microtubule cytoskeletal structures for its reproduction, multiplication or dissemination. Due to the small size of this parasite, its cytoskeleton has been primarily observable by electron microscopy. Here, we demonstrate that the nanoscale cytoskeleton organization is within reach using ultrastructure expansion microscopy (U-ExM). In developing microgametocytes, U-ExM allows to monitor the dynamic assembly of axonemes and concomitant tubulin polyglutamylation in whole cells. In the invasive merozoite and ookinete forms, U-ExM unveils the subpellicular microtubule arrays that confer cell rigidity. In ookinete, we additionally identify an apical tubulin ring above the subpellicular microtubules that colocalises with markers of the conoid in related Apicomplexa parasites. This microtubule structure was presumed to be lost in *Plasmodium* despite its crucial role in both motility and invasion in most apicomplexans. Here, U-ExM reveals that a divergent and reduced form of the conoid is actually conserved in the *Plasmodium* genus.

## Introduction

Malaria is caused by unicellular *Plasmodium* parasites that belong to the phylum Apicomplexa. More than 6,000 species have been described in this phylum^1^ including numerous human and animal pathogens such as *Toxoplasma gondii, Cryptosporidium* and *Eimeria* spp. During their lifecycle, apicomplexan parasites undergo multiple cellular differentiations into morphologically distinct forms sustaining either i) sexual reproduction, ii) asexual replication, or iii) dissemination via motility, egress and invasion of host cells^2^. Each of these forms relies on diverse microtubule cytoskeletal structures that either share general characteristics with those of other eukaryotic organisms or are unique to these parasites.

A unique cytoskeletal feature that distinguishes Apicomplexa from other eukaryotes is the apical complex. This complex is assembled in polarised invasive stages called zoites. It includes specialized secretory organelles and a microtubule-organizing centre (MTOC) called the apical polar ring (APR). The secretory organelles release various factors necessary for motility, egress or invasion. An array of subpellicular microtubules nucleates from the APR and confers the shape and stability to the cell. Two additional electron dense rings have been described above the large and thick APR in multiple apicomplexan parasites, including *Plasmodium* zoites^3,4^. The fine structure and molecular composition of the apical complex differs among apicomplexan parasites, which probably reflects different mechanical or functional requirements linked to the range of invaded host cells. For example, the presence or absence of a hollow tapered barrel structure composed of tubulin called the conoid has defined two classes within the Apicomplexa. The Conoidasida such as coccida, including *T. gondii* and gregarines, have a conoid^5^. Based on ultrastructural data, *Plasmodium* has been traditionally considered to lack a conoid and belongs to the Aconoidasida^6^. If the structure of the apical complex is relatively well characterised in some apicomplexan parasites such as *T. gondii*, its molecular composition remains elusive and difficult to visualise in *Plasmodium*.

Our understanding of *Plasmodium* microtubule structures heavily relies on electron microscopy (EM). Fluorescent light microscopy is instrumental to complement EM with the possibility of using multiple markers to infer the dynamic and the molecular composition of *Plasmodium* cytoskeletons. However, owing to the small size of the parasites, subcellular imaging by fluorescent light microscopy still poses a major challenge in *Plasmodium*. Super-resolution techniques have recently been implemented but Structured-Illumination Microscopy (SIM) provides only slightly improved resolution (~120 nm) in comparison to diffraction-limited light microscopy (~240 nm), while Stimulated Emission Depletion (STED) microscopy can reach a resolution of up to 35 nm^7,8^. However, the iron-rich hemozoin crystals present in the asexually replicating blood stages and sexual stages causes cell disintegration when illuminated with the high-power STED laser^9,10^. The implementation of guided- or rescue-STED has circumvented this issue by automatically deactivating the STED depletion laser in highly reflective regions of the sample to prevent local damage^9,10^. Because of the limit of resolution of fluorescent microscopy or the technical limitations of super resolution microscopy, the structure and molecular composition of the *Plasmodium* microtubule cytoskeleton remain difficult to interrogate.

Here, we reasoned that physical expansion of *Plasmodium* cells in an isotropic fashion by ultrastructure expansion microscopy^11^ (U-ExM) could provide an accessible bridge between traditional fluorescent microscopy and EM. We expanded multiple *Plasmodium* stages and species including two zoite stages, *P. falciparum* schizonts and *P. berghei* ookinetes, as well as developing *P. berghei* microgametes. We show that U-ExM resolves the structure of the axonemes, the mitotic hemispindles as well as the sub-pellicular microtubules. Importantly, it reveals that an apical tubulin ring colocalising with markers of the conoid in other parasites is atop the APR of motile ookinetes, suggesting that a conoid was retained in this genus. In conclusion, U-ExM enables visualization of cytoskeletal structures at a nanoscale resolution in *Plasmodium* and represents an accessible method without the need of specialized microscopes.

## Results

### U-ExM is effective in diverse stages of the *Plasmodium* lifecycle

To test the potential of U-ExM^11^ to image *Plasmodium*, we compared protocols for fluorescent microscopy of tubulin structures with or without expansion on multiple lifecycle stages (**Figure 1**). Briefly, *Plasmodium* parasites were first sedimented on poly-D-lysine coated coverslips prior to cold methanol fixation. Next, proteins of the samples were anchored to a swellable polymer, followed by denaturation and expansion. Immunostaining was subsequently performed post-expansion as previously described^11^ (see Material and Methods) (**Figure 1A**).

**Figure 1.**
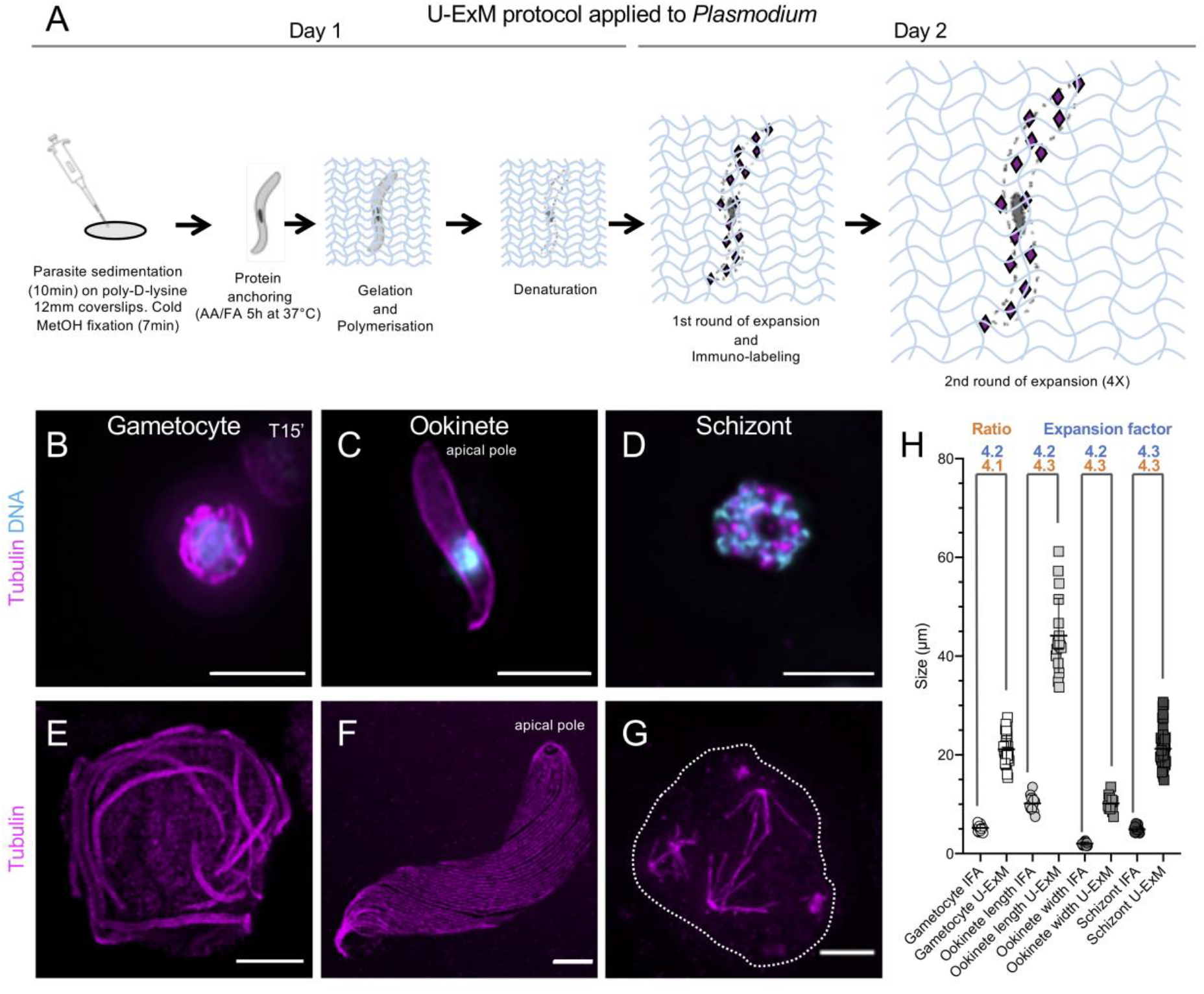
Ultrastructure expansion microscopy applied to *Plasmodium*. (**A**) Schematic illustration of ultrastructure expansion microscopy (U-ExM) protocol applied to *Plasmodium* samples. (**B-D**) Epifluorescence images of a *P. berghei* gametocyte (B), a *P. berghei* ookinete (C) and a *P. falciparum* schizont (D) stained for α- and β-tubulin (magenta, Alexa 568) and DNA (cyan). Scale bar: 5μm. (**E-G**) Same stages as in B-D expanded using U-ExM and stained for α/β-tubulin (magenta, Alexa 568). Scale bar: 5μm. (**H**) Average size of the different studied stages before (IFA) and after expansion (U-ExM). Gametocytes: IFA, 25; U-ExM, 27. Ookinetes: IFA, 15; U-ExM, 17. Schizonts: IFA, 28; U-ExM, 45.

We first imaged the development of *P. berghei* microgametocytes into microgametes, a process that is essential to establish an infection in the mosquito (**Figure 1B**). During microgametogenesis, the component parts of eight axonemes are assembled within the gametocyte cytoplasm to form flagellated male gametes in a process called exflagellation. Confocal imaging of cells stained for α/β-tubulin showed a bundle of undistinguishable axonemes coiled around the octoploid nucleus. In contrast, expanded gametocytes resolved individual axonemes (**Figure 1E**), featuring the gain of resolution obtained using U-ExM. Importantly, comparison of the average diameters of cells with or without expansion indicated a ~4.2-fold size increase in the linear dimension, from around 5 to 20 μm (**Figure 1H** and **Figure S1**).

We then applied U-ExM to the ookinete of *P. berghei*, a large motile extracellular zoite of 10 to 12 μm, responsible for colonisation of the mosquito midgut that displays a subpellicular microtubule network radiating from the apical polar ring. Non-expanded ookinetes stained for α/β-tubulin showed a diffuse cytoplasmic signal with an enrichment at the cell periphery and no visible microtubule structures (**Figure 1C**). Expanded cells showed a ~4.2-fold increase in both width and length, highlighting the isotropic expansion of the specimen (**Figure 1F, H** and **Figure S1**). This resulted in the clear resolution of individual sub-pellicular microtubules that radiate from the APR (**Figure 1F** and **Figure S1**).

Finally, we stained schizonts of *P. falciparum*. During erythrocytic schizogony, the parasite undergoes multiple rounds of replications to form 16 to 32 merozoites per schizont. Merozoites are ~1-2 μm long zoites that upon rupture of the host erythrocyte quickly reinvade erythrocytes. Confocal imaging of late stage schizonts prior to expansion showed a punctuate signal for α/β-tubulin (**Figure 1D**). Sample expansion showed a 4.3-fold increase in global size without apparent major morphological distortions and highlighted the previously described mitotic hemispindles^12^ (**Figure 1G, H** and **Figure S1**).

Altogether, expansion of multiple stages from two *Plasmodium* species indicates a consistent ~4.2-fold size increase in the linear dimension (**Figure 1H**) and demonstrates that U-ExM can be readily applied to multiple *Plasmodium* stages revealing subtle details of its internal cytoskeleton organization.

### U-ExM allows visualisation of axonemal microtubules and an associated post-translational modification in fully reconstructed gametocytes

*Plasmodium* assembles a flagellum exclusively prior to the formation of the microgamete in the midgut of the mosquito vector^13^. Microgametogenesis is an explosive development with three rounds of genome replication and endomitoses, paralleled by the assembly of eight axonemes to form eight microgametes within 10 to 20 minutes of activation^14,15^. The axonemes display the classical organisation with nine doublets of microtubules arranged in a circular pattern surrounding a central pair of singlet microtubules^16^.

Flagellum formation in *Plasmodium* is remarkably different from known models^17^. Axonemes are assembled within the cytoplasm of the microgametocyte independently of intraflagellar transport^18^ and the flagellum is formed later, after the initiation of axoneme beating. EM observations indicate that this atypical and explosive synthesis is associated with frequent axoneme miss-assembly^19^. As the exact mode of assembly remains unknown, an understanding as to the specific origins of these presumed errors might inform us of the molecular mechanisms of axoneme pattern determination^19^.

Therefore, we undertook the analysis of axonemal assembly *in cellulo* in microgametocytes using U-ExM (**Figure 2A-D**). To do so, we immunolabelled for α/β tubulin microgametocytes that were fixed 15 minutes after gametogenesis activation, when the first exflagellation events are observed. This revealed snapshots of axonemal assembly that have only been accessible by EM observations of serial sections. We observed a wide range of patterns, ranging from incomplete axoneme formation where fan-shaped arrays of short singlets or doublets microtubules radiate from the basal body (**Figure 2A** and **Figure S2**) to apparently fully assembled axonemes (**Figure 2C**). A detailed analysis of 25 whole cell projections revealed that only 4% (1/25) of 15 min activated gametocytes showed fully-formed axonemes while the others cells presented three main and non-exclusive intermediate states. First, a large number of cells (20/25) showed less than eight axonemes, which could reflect defects in basal body replication or other early defects preventing microtubule polymerisation (**Figure 2A, B** and **Figure S2**). Second, we observed that within single gametocytes, both incomplete and fully-assembled axonemes coexist within a shared cytoplasm (**Figure 2B** and **Figure S2**). Third, 44% of the imaged axonemes displayed apparently free singlets or doublets microtubules (**Figure 2B, C**, white arrows and **Figure S2**). In contrast to the frequent observation of miss-assembled axonemes in developing gametocytes, the axoneme of free microgametes did not show such apparent defects (**Figure 2D**).

**Figure 2.**
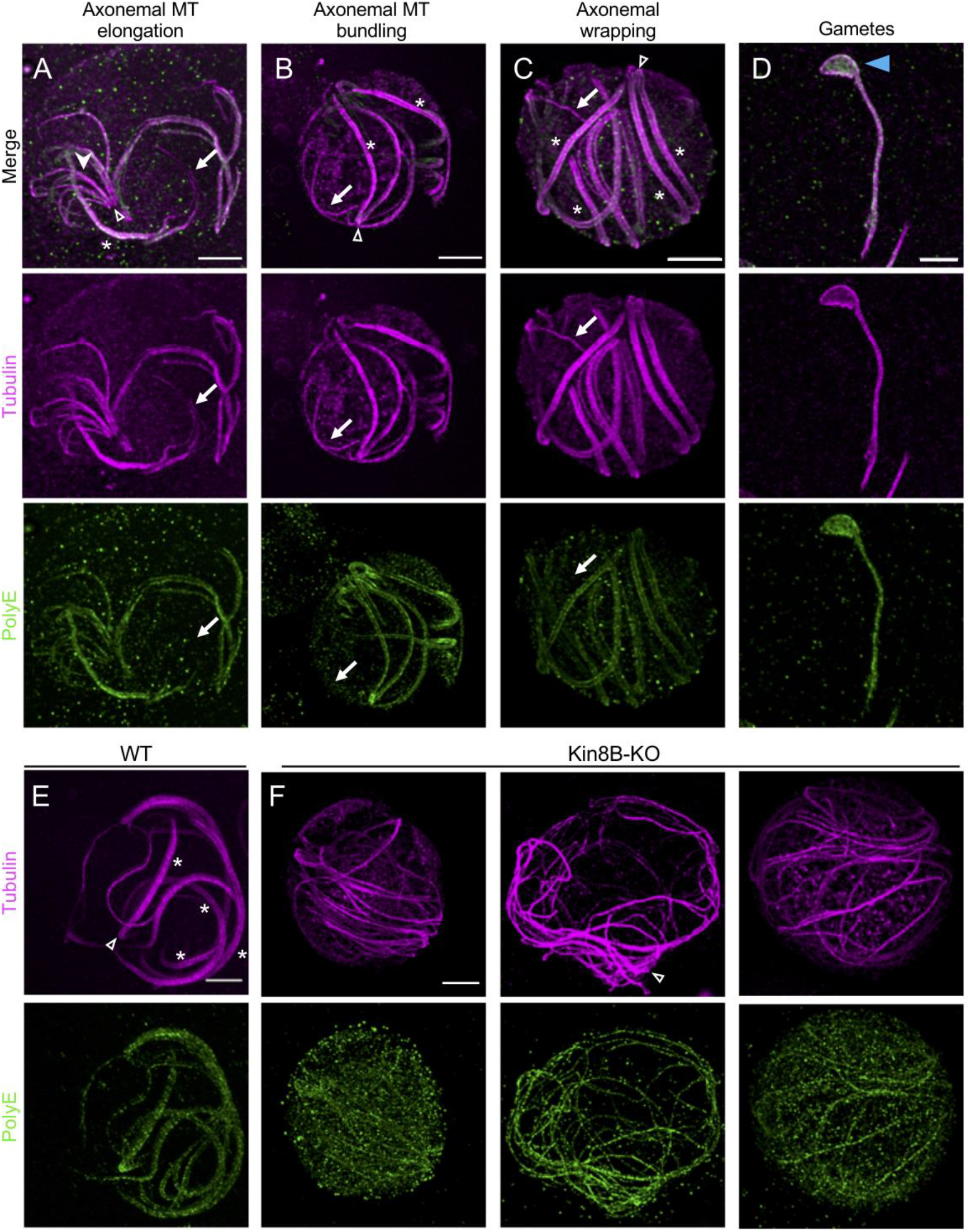
Axonemes formation in microgametocytes visualized by U-ExM. (**A-D**) Gallery of representative confocal images in gametocytes 15 minutes post-activation based on morphological features. Gametocytes were expanded and stained for α- and β-tubulin (magenta, Alexa 568) and PolyE (green, Alexa 488). Arrows show apparent differences between tubulin and PolyE staining. White arrowhead indicates the fan-shaped arrays of microtubules. Blue arrow head points to the remnant body. Open arrowhead indicates clearly identifiable basal bodies position. White asterisk denotes fully formed axonemes. Scale bar: 5μm. (**E, F**) Expanded wild-type (E) and Kin8B-KO (F) gametocytes stained for α/β-tubulin (magenta, Alexa 568) and PolyE (green, Alexa 488). Complete axonemes are visible after 15 min of activation for the wild-type, while individual unassembled singlets or doublets microtubules are seen in the mutant. Scale bar: 5μm.

To ascertain that the observed structures correspond to non-assembled microtubules, we imaged Kin8B-KO microgametocytes where full-length singlet or doublet microtubules are formed but not assembled into the 9+2 pattern^20,21^. As previously reported, we could not observe assembled axonemes in Kin8B-KO microgametocytes (27/27) 15 min after activation (**Figure 2E, F**), demonstrating that U-ExM allows visualization of singlets or doublets microtubules not incorporated into the 9+2 pattern.

To take advantage of the versatility of U-ExM, we then stained gametocytes for polyglutamylated tubulin (PolyE), a post-translational modification that stabilises microtubules with various roles in regulating flagellar motility^22^. Polyglutamylation was observed on both assembling and full-length axonemes (**Figure 2A-D**), although we also observed that some microtubules that were not incorporated into the 9+2 pattern were lacking polyglutamylation (**Figure 2A-C**, white arrows and **Figure S2**). These results suggest that polyglutamylation is added onto assembling axonemes and seems more strongly detected in fully formed axonemes. To test whether PolyE addition is dependent on axonemal formation, we stained Kin8B-KO microgametocytes for PolyE and detected a polyglutamylation signal on microtubules from non-assembled axonemes as well as a high background in the cytoplasm of the gametocytes, possibly reflecting unincorporated polyglutamylated tubulin dimers or cross-reactivity with other glutamylated proteins whose distribution is altered in the mutant. This result suggests that polyglutamylation deposition is possibly independent from axonemal formation (**Figure 2E**).

Altogether, U-ExM unveils details of axoneme formation in whole cells to levels that have only be accessible by EM. We believe that U-ExM will reveal new aspects of *Plasmodium* gametogenesis that have been overlooked by conventional fluorescence microscopy.

### In *P. falciparum* schizonts, U-ExM resolves the hemispindles and subpellicular microtubules

We then analysed expanded 40-48 hours *P. falciparum* schizonts stained for α/β tubulin (**Figure 3**). Two distinct microtubule structures were observed in this stage by U-ExM: the mitotic hemispindles and the subpellicular microtubules.

**Figure 3.**
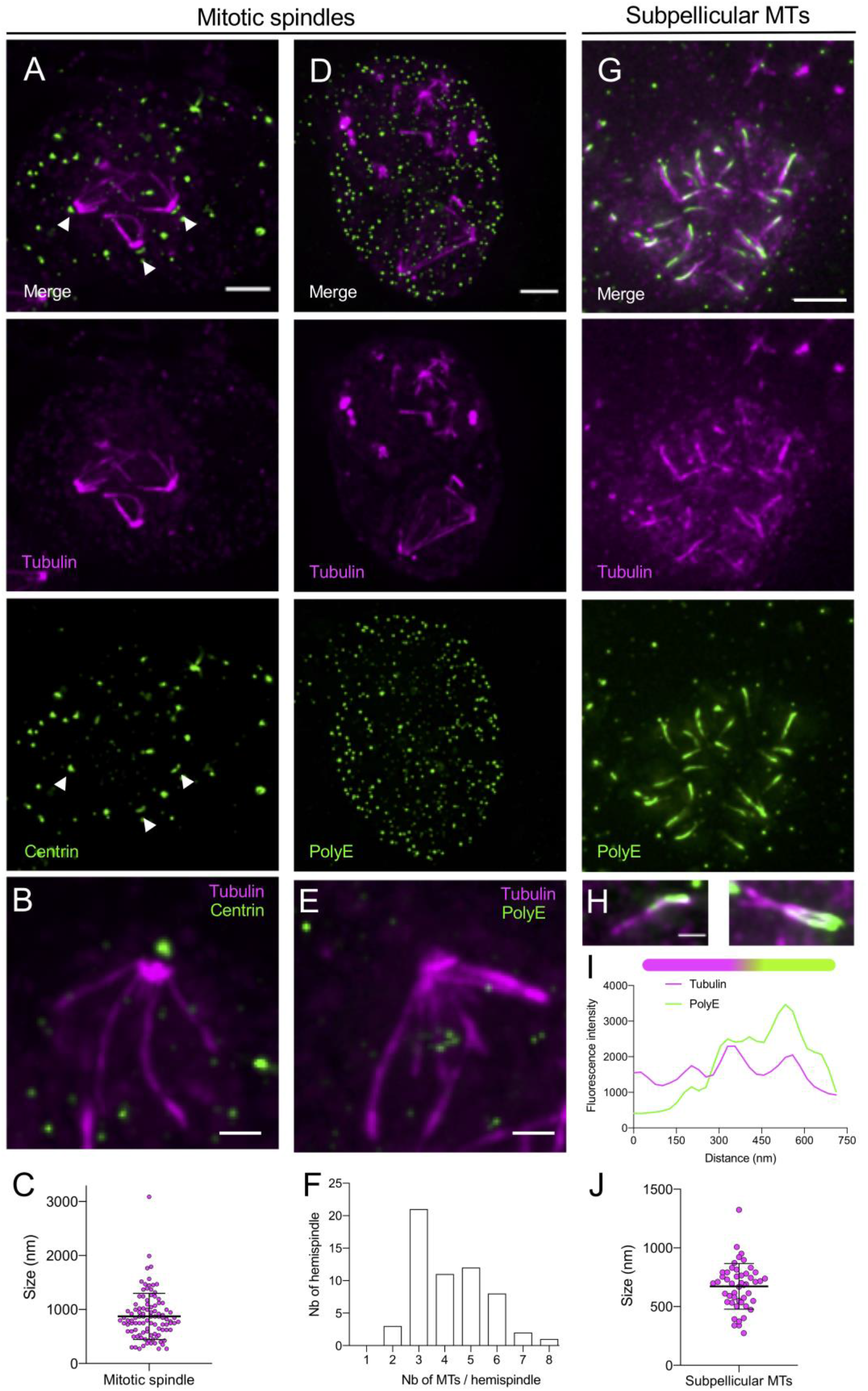
U-ExM resolves mitotic hemispindles and subpellicular microtubules in *P. falciparum* schizonts. (**A**) Epifluorescence image of an expanded schizont presenting mitotic spindles stained for α/β-tubulin (magenta, Alexa 568) and centrin (green, Alexa 488). Arrows point to centrin staining at spindle poles. Note that some extra dots are visible in the centrin staining. Scale bar: 5μm. (**B**) Zoom in of an hemispindle stained for α/β-tubulin (magenta, Alexa 568) and centrin (green, Alexa 488). Scale bar: 2μm. (**C**) Dot plot representing the mitotic spindle length. Average length +/- standard deviation: 872.68 nm +/- 427, n=102. (**D**) Epifluorescence image of an expanded schizont presenting mitotic spindles stained for α/β-tubulin (magenta, Alexa 568) and PolyE (green, Alexa 488). Scale bar: 5μm. Note that the mitotic spindle is not polyglutamylated. (**E**) Zoom in of an hemispindle stained for α/β-tubulin (magenta, Alexa 568) and PolyE (green, Alexa 488). Scale bar: 2μm. (**F**) Histogram representing the number of hemispindles displaying 1, 2, 3, 4, 5, 6, 7, or 8 microtubules. (**G**) Representative epifluorescence images of expanded schizonts presenting subpellicular microtubules stained for α/β tubulin (magenta, Alexa 568) and PolyE (green, Alexa 488). Note that the subpellicular microtubules are polyglutamylated in contrast to the mitotic spindle. Scale bar: 5μm. (**H**) Two examples of representative subpellicular microtubules stained for α/β tubulin (magenta, Alexa 568) and PolyE (green, Alexa 488). Scale bar: 5μm. (**I**) Plot profile along a subpellicular microtubule displaying tubulin and PolyE signals with a schematic representation of a subpellicular microtubule on top. Magenta: tubulin, Green: PolyE. Note that PolyE is not uniformly distributed along the subpellicular microtubules. (**J**) Dot plot representing the size of subpellicular microtubules. Average length+/- standard deviation: 672 +/- 192 nm, n=48. From 3 independent experiments.

In late schizonts, we observed the hemispindles that form an array of microtubules, which radiate from single MTOCs, as revealed by centrin staining (**Figure 3A-B** arrow heads indicate centrin dots at the mitotic spindle pole). Per centriolar plaque, we observed an average of four hemispindles ranging from 0.5 to 2 μm (**Figure 3C**), as previously described^23^. Each hemispindle contained on average three microtubules (**Figure 3F**). Interestingly, the hemispindles were not polyglutamylated as opposed to mitotic spindles in other eukaryotic cells^24^ possibly suggesting a requirement for dynamic spindles in *Plasmodium* (**Figure 3E**).

In segmenter schizonts, U-ExM resolved one to two subpellicular microtubules per merozoite that were on average 0.7 μm long (**Figure 3G-J**), as previously described by EM^3,4^ or STED microscopy^12^. As opposed to the hemispindles, subpellicular microtubules were polyglutamylated with a polar distribution of polyglutamylation (**Figure 3I-H**).

### The *P. berghei* ookinete expresses a conoid

The apical complex of *Plasmodium* zoites was shown to contain three electron dense concentric rings (known as polar rings), with the smallest directly adjacent to the apical tip^3,4^. The third proximal ring, the apical polar ring (APR) is much thicker than the two other apical rings and serves as a MTOC for the 2 to 3 subpellicular microtubules. Despite previous observations of the apical complex by ultrastructural studies, there is relatively limited knowledge of its molecular composition and variation across the different zoite stages in *Plasmodium*^25^. We thus analysed expanded ookinetes similarly stained for α/β tubulin and polyglutamylated tubulin (**Figure 4A**).

**Figure 4.**
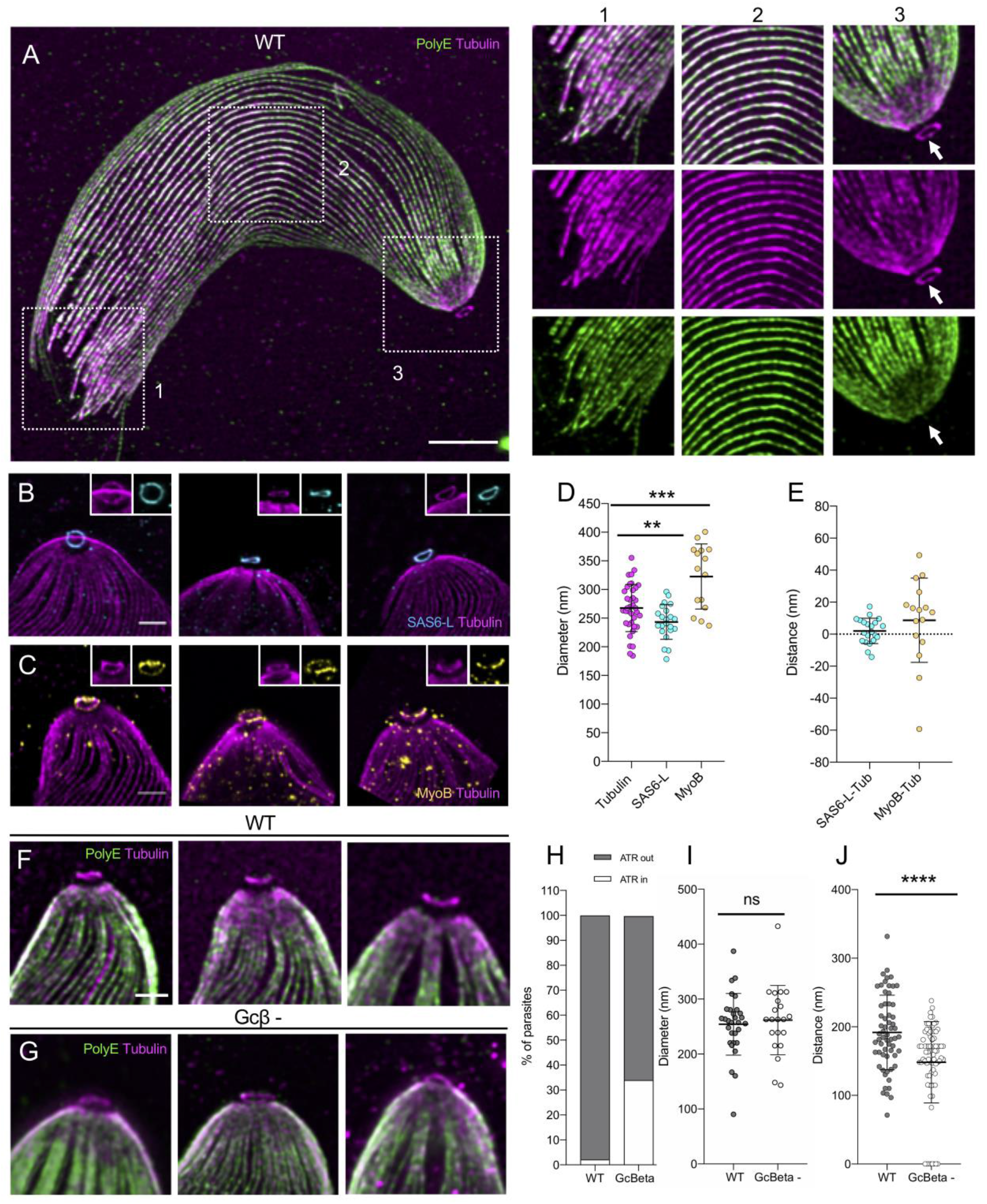
Identification and characterisation of a conoid-like structure in *P. berghei* ookinetes. **(A)** Representative confocal image of expanded ookinete stained for α/β-tubulin (magenta, Alexa 568) and PolyE (green, Alexa 488) highlighting 3 subregions boxed in white. 1: distal, 2: centre and 3: apical regions, respectively. Note that the tubulin ring or ATR in 3 is not polyglutamylated. Scale bar: 5μm. (**B**) Gallery of the apical region of expanded ookinetes stained for α/β-tubulin (magenta, Alexa 568) and SAS6L (cyan, Alexa 488). Scale bar: 1μm. (**C**) Gallery of the apical region of expanded ookinetes stained for α- and β-tubulin (magenta, Alexa 568) and MyoB (yellow, Alexa 488). Scale bar: 1μm. (**D**) Measure of the diameter of the ATR (Tubulin), SAS6L (cyan) and MyoB (yellow) rings. Averages and standard deviations are as follows: Tubulin: 237.69 +/- 41.11 (n=38), SAS6L: 243.25 +/- 30.22 (n=24), MyoB: 332.64 +/-56.72 (n=15) nm. Data from 2 independent experiments. Student t test: p = 0.0186 (SAS6L versus Tubulin) and p = 0.0003 (MyoB versus Tubulin). (**E**) Distance between the tubulin rings and SAS6L (cyan) or MyoB (yellow). Averages and standard deviations are as follows: SAS6L-Tub: 2.01 +/- 7.88 (n=20) and MyoB-Tub: 8.70 +/- 26.38 (n=15) nm. Data from 2 independent experiments. (**F, G**) Gallery of the apical region of expanded wild-type (F) and GCβ^-^ mutant (G) ookinetes stained for α/β-tubulin (magenta, Alexa 568) and PolyE (green, Alexa 488). Scale bar: 1 μm. (**H**) Percentage of wild type and GCβ^-^ mutant ookinetes displaying a visible or collapsed ATR. N=40 from 3 independent experiments. (**I**) Measurement of the ATR diameter in wild type and GCβ^-^ expanded ookinetes. Averages and standard deviations are as follows: WT= 254.14 +/- 56.18 and GCβ^-^= 261.73 +/- 63.05 nm. n= 27 from 3 independent experiments. We observed no difference in the measured diameters in the two conditions. Student t test: p= 0.6534. ns= not significant. (**J**) Distance between the apical pole to the ATR in wild type and GCβ^-^ mutant expanded ookinetes. Averages and standard deviations are as follows: WT= 191.81 +/- 54.31 and GCβ^-^= 163.08 +/- 32.26 nm. N= 61 from 3 independent experiments. Student t test: p< 0.0001.

A significantly larger number of sub-pellicular microtubules (>40) that radiate from the APR and covering most of the ookinete body were distinctly observed in expanded ookinetes. As in segmented schizonts, the subpellicular microtubules were polyglutamylated (**Figure 4A** and **Figure S3**). A less intense signal for polyglutamylation was observed at the apical extremity of the ookinete likely corresponding to the electron dense collar structure^26^ (**Figure 4A** and **Figure S3**).

Importantly, the improved resolution of U-ExM in ookinetes allowed us to observe a ring of tubulin above the apical polar ring that we named the Apical Tubulin Ring (ATR) (**Figure 4A**), a feature that was never described before. This was surprising as such a ring of tubulin is reminiscent of a conoid found in coccidian but thought to be lost in the Haemosporidia order containing *Plasmodium*^2^. In *T. gondii* tachyzoites, the conoid is a central component of the apical complex and consists in a set of tubulin filaments that create a cone-shaped structure located between the APR and two additional apical rings called the pre-conoidal rings^27^. A pair of intra-conoidal microtubules additionally traverses the pre-conoidal rings and the conoid. We thus wondered whether the ATR could represent a structure related to the conoid in *Plasmodium*.

First, we noticed that the ATR is not polyglutamylated as previously described for the conoid of *T. gondii*^2^ (**Figure 4A**). While the conoid is thought to be absent in the *Plasmodium* lineage, some proteins associated to the conoid in *Toxoplasma* are also conserved in *Plasmodium*^25^. This includes the SAS-6-like (SAS6L) protein that was shown to localise to the conoid in *Toxoplasma*^29^ and that has been demonstrated to form an apical ring in *Plasmodium* ookinetes^30^. In expanded ookinetes, endogenously tagged SAS6L-GFP^30^ co-localized with the ATR (**Figure 4B**). While the distance between the SAS6L-GFP/ATR ring and the APR was the same, the diameter of the SAS6L-GFP ring was slightly shorter suggesting that SAS6L-GFP or at least its C-terminus is facing the interior of the ATR (**Figure 4D, E**). In *Plasmodium*, MyoB was also shown to localise at a discrete apical location in merozoite and ookinete^31^ and was suggested to fulfil, at least in part, a similar role as the conoid-associated myosin H in *T. gondii*^32^. U-ExM revealed that endogenously tagged MyoB-GFP^31^ formed a dotty ring at the ookinete apex (**Figure 4C**). However, this ring was larger and slightly above the ATR/SAS6L-GFP ring, further suggesting the presence of multiple apical rings in ookinetes (**Figure 4C-E**). However, despite the presence of the ATR, no intra-conoidal microtubule could be observed.

Another characteristic of the *T. gondii* conoid lies in its ability to protrude through the apical polar ring upon stimulation of secretion and motility^33^. We thus tested whether the ATR position was linked to the motility of ookinete by comparing its position in motile and non-motile ookinetes (**Figure 4F-J**). Ookinetes rely on the cGMP-dependent protein kinase G (PKG) to sustain productive secretion and gliding motility^34,35^. To determine the position of the ATR upon PKG activation in wild-type parasites, we imaged mutants of the cGMP-producing guanylyl cyclase beta (GCβ) that show severely reduced motility, with only rare bouts of slow gliding^34,36,37^. In motile wild-type ookinetes, the distance between the APR and the ATR was of 200 nm while in 34% of GCβ^-^ ookinetes, the ATR was in close proximity to the APR (**Figure 4H**). The remaining mutant cells showed a significantly reduced distance of 160 nm between the ATR and the APR (**Figure 4J** and **Figure S4**). These results suggest that the position of the ATR relative to the APR depends on the activation of secretion and motility in ookinetes.

## Discussion

Our results demonstrate that U-ExM can be applied successfully to several *Plasmodium* species and lifecycle stages to unveil ultrastructural details of the microtubule cytoskeleton that was, up to now, only reachable by electron or super resolution microscopy. Imaging forming axonemes by U-ExM during microgametogenesis gave new insights into their mode of assembly (**Figure 5A**). For example, we show that post-translational modifications such as polyglutamylation occur in forming axonemes. We also observed different and non-exclusive states with either less than eight axonemes per microgametocyte, axonemes with different lengths within a cell or frequent mis-incorporation of microtubules toward the distal extremity of the axoneme. It remains unknown whether these observations reflect transient intermediate stages of assembly or terminal errors, as previously suggested^19^. However, as free gametes did not show obvious defects in axoneme assembly, it is tempting to speculate that microtubules may still grow and bundle upon the initiation of beating. Another possibility would be that only fully-formed mature axonemes initiate efficient beating to form motile microgametes even in the presence of other defective axonemes in the same microgametocyte (**Figure S2**). Further experimental characterisations will be required to test these hypotheses.

**Figure 5.**
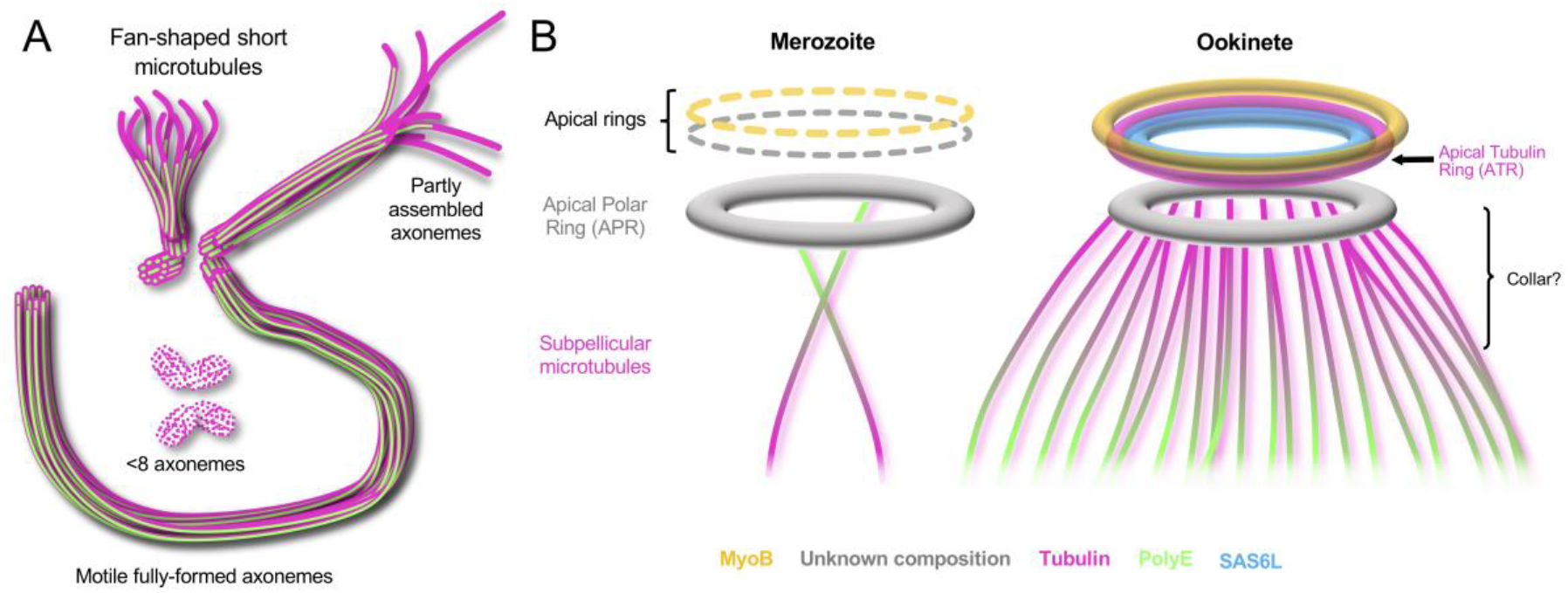
Current models. **(A)** Proposed model of axonemal assembly in microgametocytes featuring the co-existence of short fan-shaped microtubules with partly assembled and fully assembled axonemes. Basal bodies are represented at the base of axonemes. Microtubules: magenta, and PolyE: green. **(B)** Proposed model of the apical complex organisation in *Plasmodium* merozoite and ookinete highlighting a conoid-like structure present in ookinete, composed of an apical tubulin ring (ATR, magenta) and SAS6L (cyan) above the apical polar ring (grey). An additional MyoB positive ring is detected in both merozoite and ookinete (yellow). Grey denotes the presence of additional electron dense rings observed by electron microscopy whose molecular composition remains unknown. The relative position of MyoB in merozoite is currently unknown. The orientation of the polyglutamylation profile of the subpellicular microtubules in the merozoite is proposed by analogy with the polyglutamylation profile of subpellicular microtubules in *T. gondii* tachyzoites^28^. In ookinete the apical section of less polyglutamylated subpellicular microtubules may correspond to the collar.

The most striking feature revealed by U-ExM was the presence of an ATR in ookinetes that most likely corresponds to a *Plasmodium* conoid. Even though the structure of the ATR appears significantly reduced or compacted compared with the spiralling tubulin-rich fibres that form the conoid in other members of Apicomplexa, multiple lines of evidence indicate that the ATR is related to the conoid (**Figure 5B**). First, the ATR co-localises closely with the SAS6L ring^30^ that was previously shown to be associated with the conoid in *T. gondii*^29^. Second, the ATR position relative to the APR depends on the activation of ookinete motility and secretion, a characteristic reminiscent of the enigmatic protrusion of the conoid that happens upon activation of secretion and motility in *T. gondii* tachyzoites^33^. Third, the ATR is associated with at least one additional tightly apposed ring that may correspond to one of the so-called preconoidal rings in *T. gondii*. This additional ring is composed of MyoB, which was proposed to fulfil, at least in part, a similar role as the conoid-associated myosin H in *T. gondii*^22^. Interestingly, while MyoB is also expressed in schizonts^31^, the ATR/SAS6L ring is not observed at this stage^30^, further confirming that the *Plasmodium* apical complex shows stage-specific variations in its composition, as previously proposed^25,30^. In addition to SAS6L and MyoB, multiple other proteins associated with the conoid in *T. gondii* are expressed in *Plasmodium*^25,33^ and we expect that U-ExM will allow to define their exact position in the *Plasmodium* apical complex.

We thus propose that *Plasmodium* parasites retained a conoid that is significantly divergent and reduced compared with the best-described *T. gondii* conoid, which likely prevented its identification. Our results also suggest that the tubulin structure that initially defined the conoid is not expressed in all *Plasmodium* zoites raising new questions on the molecular nature of the conoid. It now appears that the conoid is likely a conserved organelle in most Apicomplexa parasites including *Plasmodium*. However, its molecular composition is more diverse that initially expected and the exact physiological requirements associated with this diversity remain mysterious. It was suggested that the conoid plays a mechanical role in the invasion by parasites such as *Toxoplasma* that must penetrate the robust barrier of the intestinal epithelium of vertebrates^2^. The observation variations in the molecular composition of the conoid between schizont and ookinete thus likely reflect adaptations of the apical complex to the different host cells or environment encountered by these two zoites.

Altogether, this work highlights the potential of using expansion microscopy to reveal the microtubule network in diverse stages of *Plasmodium* species with unprecedented resolution. Moreover, expansion microscopy methods have been successfully applied with super-resolution approaches, leading to a further increase in resolution^11^. Such approaches might pave the way for further investigation of the *Plasmodium* cytoskeleton and its molecular organization. Finally, the recently developed membrane-based expansion microscopy protocols^38,39^ could also provide other exciting insights into the membrane biology of *Plasmodium*. In summary, the use of expansion microscopy can nicely complement electron and super resolution microscopy as it provides molecular details to the available structural information without the need for specialised microscopes.

## Author Contributions

V.H, P.G. and M.B. conceived, supervised and designed the project. E.B and L.B performed the experiments and analysed the data. A.B. provided all *Plasmodium* samples as well as the associated expertise. All authors wrote the manuscript.

## Acknowledgments

We thank the BioImaging Center at Unige as well as Philippe Bastin, Linda Kohl, Dominique Soldati-Favre, Nicolas Dos Santos Pacheco, Julien Guizetti, and Nikolai Klena for critical reading of the manuscript and insightful comments. We also would like to thank Rita Tewari for generously sharing the *P. berghei* Kin8B-KO, SAS6L-GFP and MyoB-GFP lines. This work was supported by the Swiss National Science Foundation (SNSF) PP00P3_187198 attributed to P.G, BSSGI0_155852 and 31003A_179321 to MB, and by the European Research Council ERC ACCENT StG 715289 attributed to P.G. MB is an INSERM investigator. MB and PG are EMBO young investigators. EB is supported by an EMBO long-term fellowship (ALTF-284-2019).

## Data and materials availability

All data needed to evaluate the conclusions in the paper are present in the paper and/or the supplementary materials. Additional data available from authors upon request. Correspondence and requests for materials should be addressed to V.H., P.G. or M.B.

## Supplementary material

**Figure S1.**
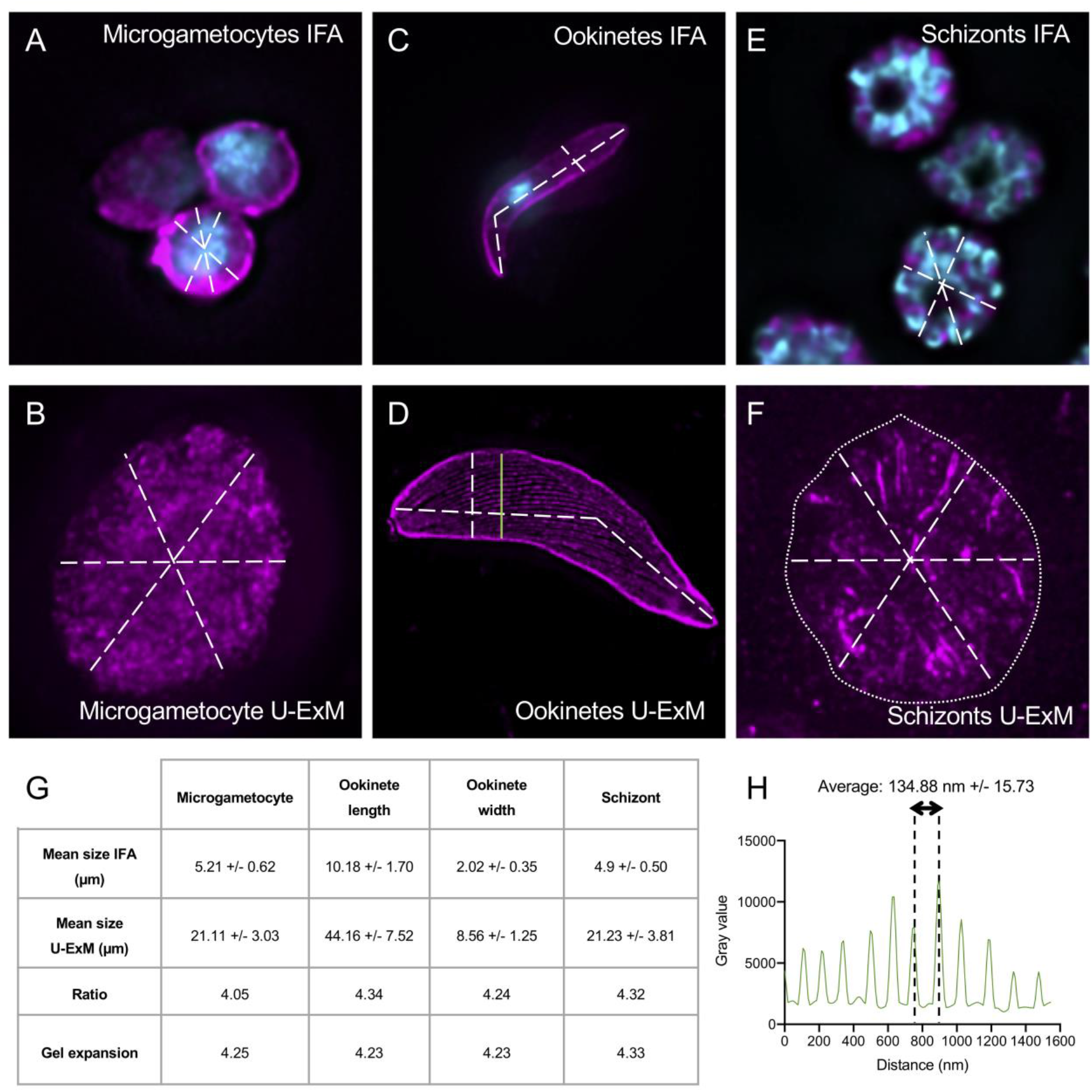
Measurements to assess the isotropicity of U-ExM expanded specimens. Measurement skills to evaluate parasite size before and after expansion. Dotted lines represented ROI used for the measurements for each parasite stage: microgametocytes, nonexpanded (**A**) and expanded (**B**); ookinetes non-expanded (**C**), and expanded (**D**); schizonts non-expanded (**E**) and expanded (**F**). Scale bar: 5μm. (**G**) Ratio between the size before and after expansion and the average expansion factor (gel size/coverslip size). Averages and standard deviations are given in the table from at least 3 independent experiments for each stage. Microgametocytes n=25, ookinete length n=27, ookinete width=14, and schizonts n=30. (**H**) Plot profile of the region of interest extracted from one ookinete image stained for α/β-tubulin (magenta, Alexa 568) showing the resolution limit of U-ExM. The example of ROI used for this plot profile is indicated in green on panel D. The average distance between two microtubules is of 135 nm +/- 16.

**Figure S2.**
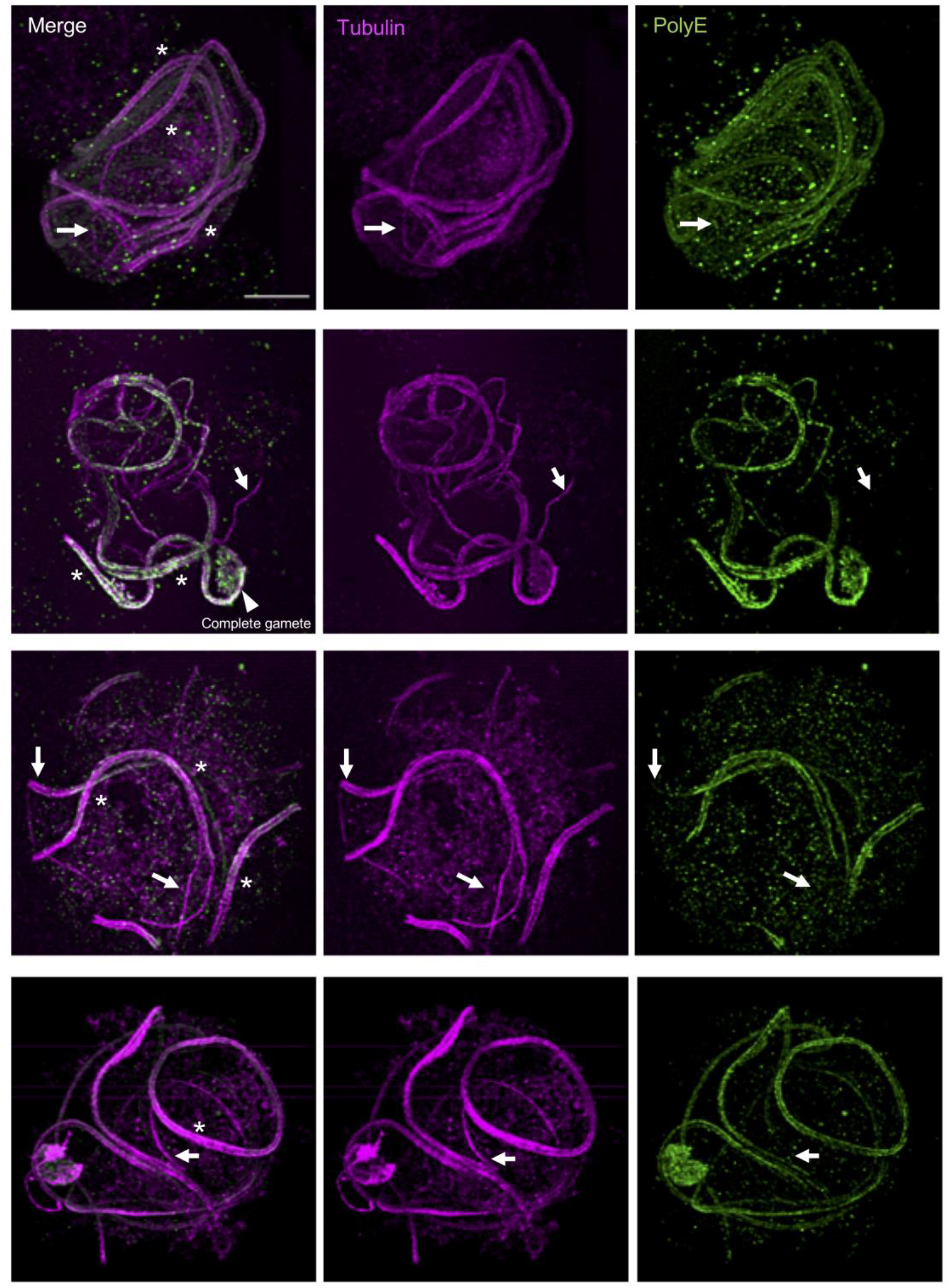
Gallery of U-ExM expanded wild-type *Plasmodium* microgametocytes. Expanded WT microgametocytes and microgametes stained for α- and β-tubulin (magenta, Alexa 568) and PolyE (green, Alexa 488). Arrows show apparent differences between tubulin and PolyE staining. White asterisk denotes fully formed axonemes. The white arrow head indicates apparently exflagellated gamete. Scale bar: 5 μm.

**Figure S3.**
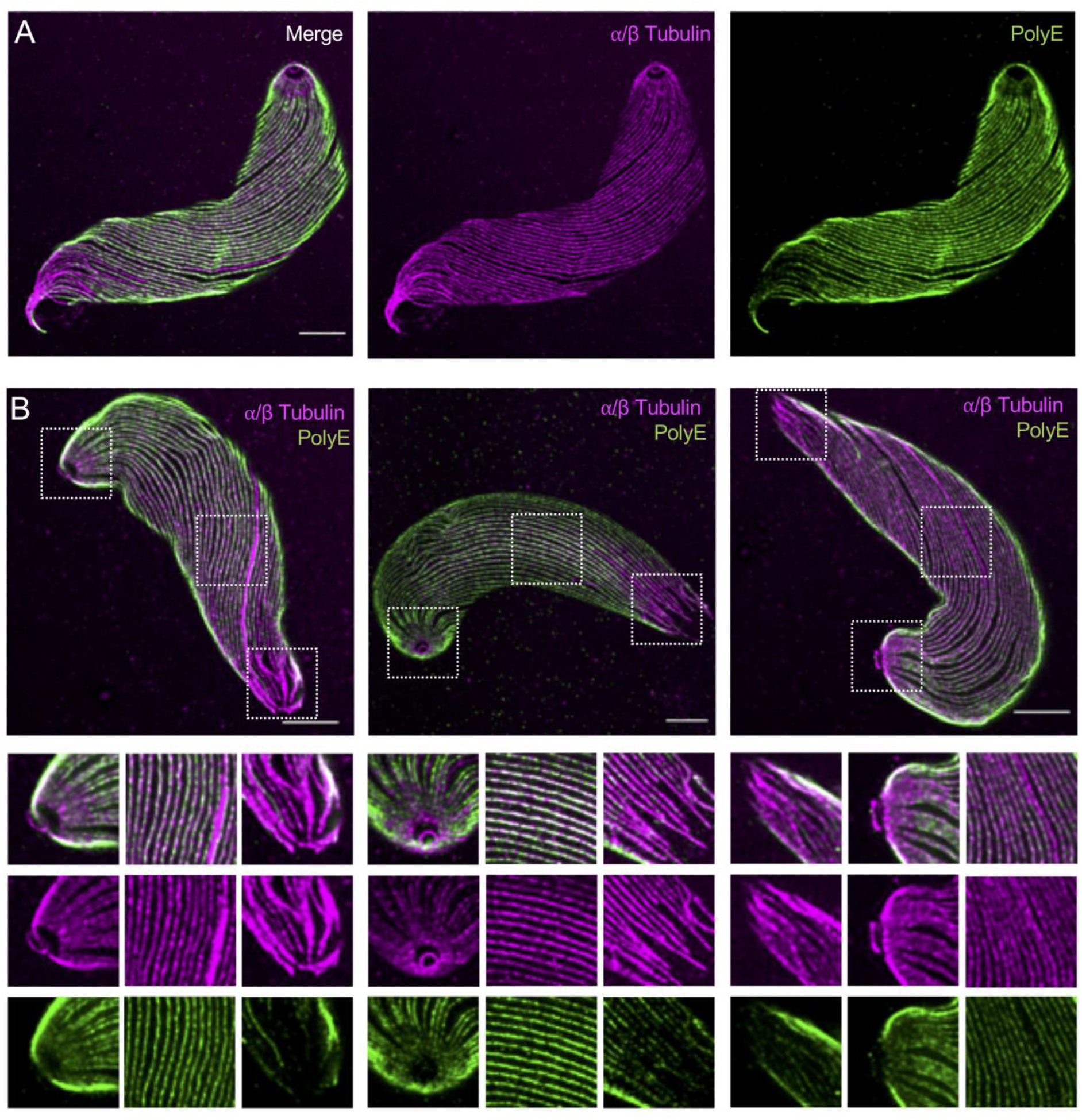
U-ExM of wild-type *Plasmodium* ookinetes. (**A**) Representative confocal image of wild type ookinete expanded using the U-ExM protocol and stained for α/β-tubulin (magenta, Alexa 568) and PolyE (green, Alexa 488). The two channels are represented independently. Scale bar: 5μm. (**B**) Representative gallery of confocal images of wild-type ookinetes. Ookinetes were expanded using the U-ExM protocol and stained for α/β-tubulin (magenta, Alexa 568) and PolyE (green, Alexa 488). Scale bar: 5μm. Insets show the delineated white-boxed region. Note the apical tubulin ring (ATR) is not polyglutamylated and the subpellicular microtubules that are PolyE positive.

**Figure S4.**
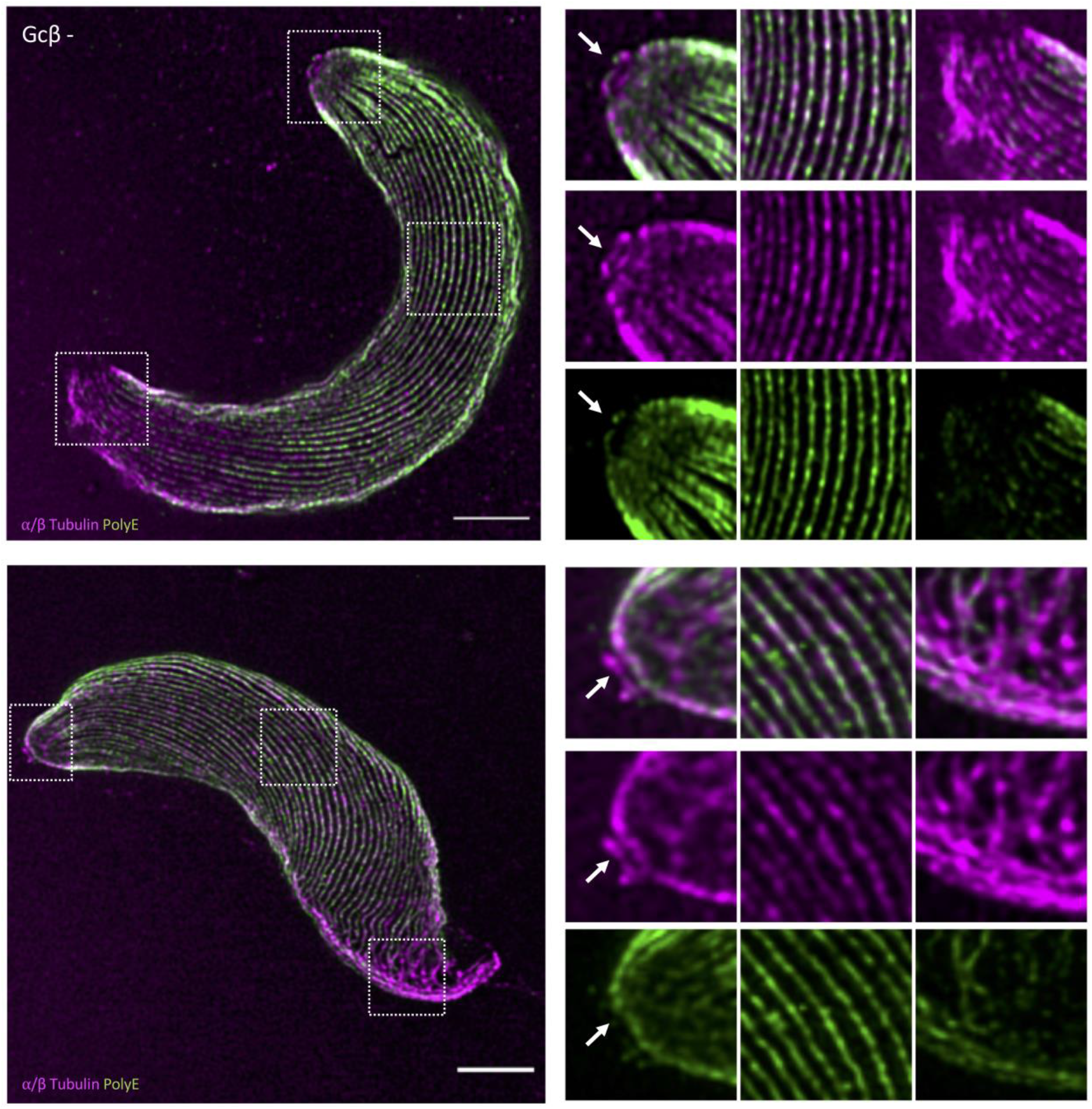
U-ExM of expanded GCβ^-^ *Plasmodium* ookinetes. Representative confocal images of GCβ^-^ ookinete mutants. Ookinetes were expanded using the U-ExM protocol and stained for α/β-tubulin (magenta, Alexa 568) and PolyE (green, Alexa 488). Scale bar: 5μm. The ATR is closely associated with the apical polar ring in contrast to wild-type ookinetes.

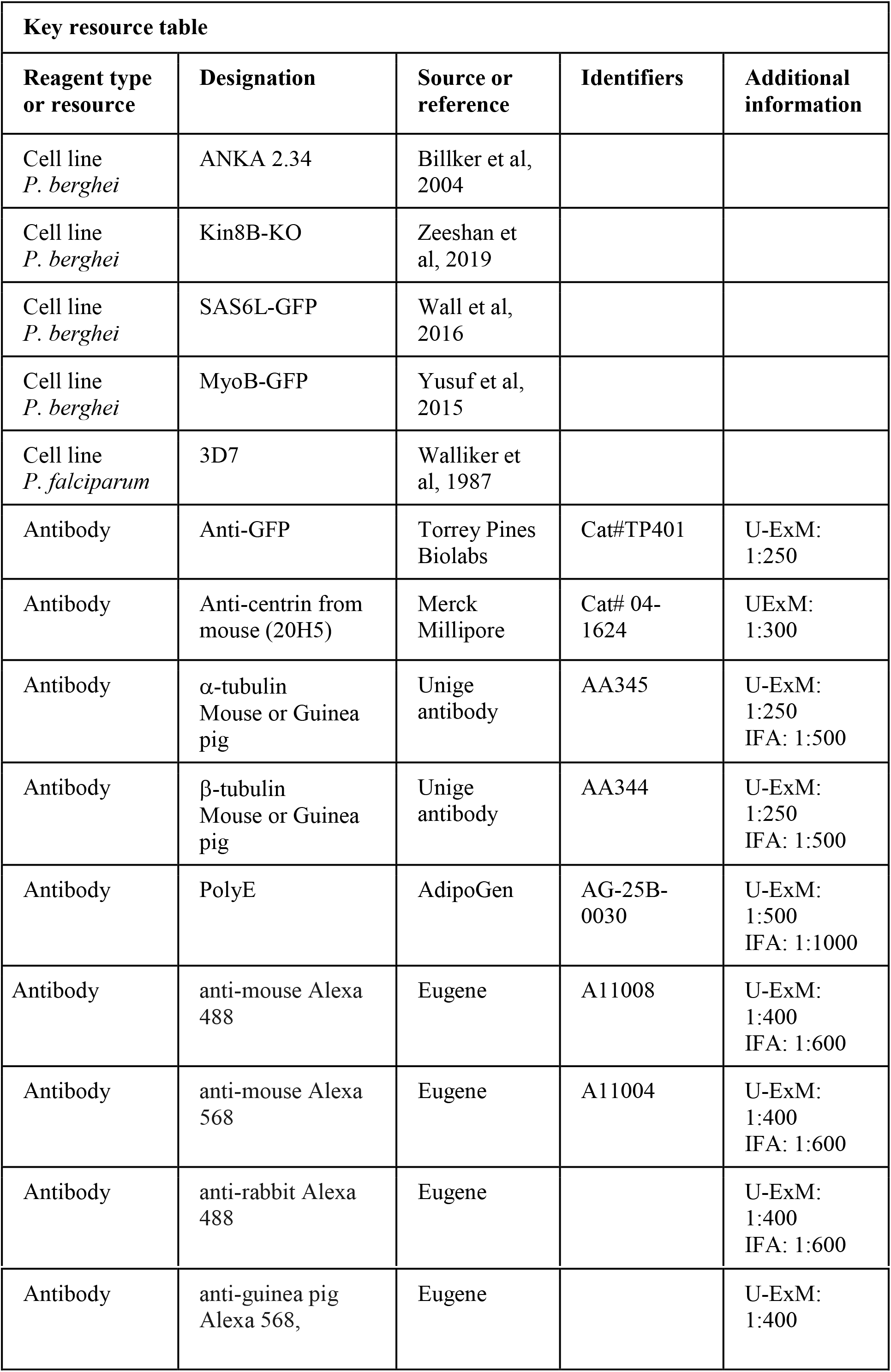

| Key resource table |  |  |  |  |
| --- | --- | --- | --- | --- |
| Reagent type or resource | Designation | Source or reference | Identifiers | Additional information |
| Cell line<br>*P. berghei* | ANKA 2.34 | Billker et al, 2004 |  |  |
| Cell line<br>*P. berghei* | Kin8B-KO | Zeeshan et al, 2019 |  |  |
| Cell line<br>*P. berghei* | SAS6L-GFP | Wall et al, 2016 |  |  |
| Cell line<br>*P. berghei* | MyoB-GFP | Yusuf et al, 2015 |  |  |
| Cell line<br>*P. falciparum* | 3D7 | Walliker et al, 1987 |  |  |
| Antibody | Anti-GFP | Torrey Pines Biolabs | Cat#TP401 | U-ExM: 1:250 |
| Antibody | Anti-centrin from mouse (20H5) | Merck Millipore | Cat# 04-1624 | UExM: 1:300 |
| Antibody | α-tubulin<br>Mouse or Guinea pig | Unige antibody | AA345 | U-ExM: 1:250<br>IFA: 1:500 |
| Antibody | β-tubulin<br>Mouse or Guinea pig | Unige antibody | AA344 | U-ExM: 1:250<br>IFA: 1:500 |
| Antibody | PolyE | AdipoGen | AG-25B-0030 | U-ExM: 1:500<br>IFA: 1:1000 |
| Antibody | anti-mouse Alexa 488 | Eugene | A11008 | U-ExM: 1:400<br>IFA: 1:600 |
| Antibody | anti-mouse Alexa 568 | Eugene | A11004 | U-ExM: 1:400<br>IFA: 1:600 |
| Antibody | anti-rabbit Alexa 488 | Eugene |  | U-ExM: 1:400<br>IFA: 1:600 |
| Antibody | anti-guinea pig<br>Alexa 568, | Eugene |  | U-ExM:<br>1:400 |

## Materials and Methods

### Ethics statement

All animal experiments were conducted with the authorisation numbers GE/82/15 and GE/41/17, according to the guidelines and regulations issued by the Swiss Federal Veterinary Office.

### *P. berghei* maintenance and production

*P. berghei* ANKA strain^40^-derived clone 2.34^41^ together with derived transgenic lines Kin8B-KO^20^, SAS6L-GFP^30^, MyoB-GFP^31^ were grown and maintained in CD1 outbred mice as previously described^42^. Six to ten week-old mice were obtained from Charles River laboratories, and females were used for all experiments. Mice were specific pathogen free (including *Mycoplasma pulmonis*) and subjected to regular pathogen monitoring by sentinel screening. They were housed in individually ventilated cages furnished with a cardboard mouse house and Nestlet, maintained at 21 ± 2 °C under a 12 hr light/dark cycle, and given commercially prepared autoclaved dry rodent diet and water *ad libitum*. The parasitaemia of infected animals was determined by microscopy of methanol-fixed and Giemsa-stained thin blood smears.

Gametocyte production and purification was performed as previously described^43^. Parasites were grown in mice that had been phenyl hydrazine-treated three days before infection. One day after infection, sulfadiazine (20 mg/L) was added in the drinking water to eliminate asexually replicating parasites. Microgametocyte exflagellation was measured three or four days after infection by adding 4 μl of blood from a superficial tail vein to 70 μl exflagellation medium (RPMI 1640 containing 25 mM HEPES, 4 mM sodium bicarbonate, 5% fetal calf serum (FCS), 100 μM xanthurenic acid, pH 7.4). To calculate the number of exflagellation centres per 100 microgametocytes, the percentage of red blood cells (RBCs) infected with microgametocytes was assessed on Giemsa-stained smears. For gametocyte purification, parasites were harvested in suspended animation medium (SA; RPMI 1640 containing 25 mM HEPES, 5% FCS, 4 mM sodium bicarbonate, pH 7.20) and separated from uninfected erythrocytes on a Histodenz cushion made from 48% of a Histodenz stock (27.6% [w/v] Histodenz [Sigma/Alere Technologies] in 5.0 mM TrisHCl, 3.0 mM KCl, 0.3 mM EDTA, pH 7.20) and 52% SA, final pH 7.2. Gametocytes were harvested from the interface.

Ookinete cultures were performed as previously described^35^. Parasites were maintained in phenyl hydrazine-treated mice. Ookinetes were produced *in vitro* by adding 1 volume of high gametocytaemia blood in 30 volumes of ookinete medium (RPMI1640 containing 25 mM HEPES, 10% FCS, 100 μM xanthurenic acid, pH 7.5) and incubated at 19°C for 18-24 hrs. Ookinetes were purified using paramagnetic anti-mouse IgG beads (Life Technologies) coated with anti-p28 mouse monoclonal antibody (13.1).

### *P. falciparum* culture

*P. falciparum* strain 3D7, a clone from the NF54 isolate^44^, was grown in human erythrocytes in RPMI-1640 medium with glutamine (Gibco), 0.2% sodium bicarbonate, 25 mM HEPES, 0.2% glucose, 5% human serum, and 0.1% Albumax II (Life Technologies). Parasite cultures were kept synchronized by double sorbitol treatments. Late stage parasites were purified from synchronous cultures using Percoll (GE Healthcare).

### U-ExM

Ookinetes and schizonts were centrifuged at 1000 rpm during 5 minutes at 24°C and resuspended in 500 μL of PBS 1X. Parasites were sedimented on poly-D-lysine (A-003-E, SIGMA) coverslips (150 μL/coverslip) during 10 minutes at RT. To stop the activation of the gametocytes, the same protocol was used but keeping the parasites at 4°C. Then parasites were fixed in −20°C methanol during 7 minutes and prepared for Ultrastructural Expansion Microscopy (U-ExM) as previously published^11^. Briefly, coverslips were incubated for 5 hours in 2X 1.4 % AA/ 2% FA mix at 37°C prior gelation in APS/ Temed /Monomer solution (19% Sodium Acrylate; 10% AA; 0,1% BIS-AA in PBS 10X) during 1 hour at 37°C. Then denaturation was performed during 1h30 at 95°C^45^. After denaturation, gels were incubated in ddH2O at RT during 30min. Next, gels were incubated in ddH2O overnight for complete expansion. The following day, gels were washed in PBS twice for 15 min to remove excess water before incubation with primary antibody solution. They were stained 3 hours at 37°C with primary antibodies against PolyE (1:500), α-tubulin and β-tubulin (1:200), centrin (1:300), anti-GFP (1:250). Gel were washed 3 x 10 minutes in PBS-Tween 0.1% prior incubation with secondary antibodies (anti-mouse Alexa 568, Anti-mouse 488, anti-rabbit Alexa 488, Anti-guinea pig Alexa 568 - 1:400) during 3 hours at 37°C and 3 washes of 10 minutes in PBS-Tween. Overnight a second round of expansion was done in water before imaging.

Imaging was performed on a Leica Thunder inverted microscope using 63X 1.4NA oil objective with Small Volume Computational Clearing mode to obtain deconvolved images. 3D stacks were acquired with 0.21 μm z-interval and x,y pixel size of 105 nm. Images were analysed and merged using ImageJ software.

Confocal microscopy was performed on a Leica TCS SP8 with a 63×/1.4-NA (numerical aperture) oil-immersion objective, using the HyVolution mode20 to generate deconvolved images, with the following parameters: ‘HyVolution Grade’ at max resolution, Huygens Essential as ‘Approach’, water as ‘Mounting Medium’, and Best Resolution as ‘Strategy’.

### Measurements

The diameter and distance between the ring of tubulin and the microtubules were always measured from dual staining experiments. After applying a maximum projection of the stack, we used the home-made plugin pickCentrioleDim on Fiji software developed by M. Le Guennec to measure the peak-to-peak distance for each fluorescent channel for each parasite. The expansion factor of each gel was after applied to the value to obtain the real distance.

To measure the length of microtubules we used Fiji to draw a line to define our ROI following tubulin staining and determined its associated length. The expansion factor of each gel was after applied to the value to obtain the real size.

